# Direct detection of CRISPR mutations and transcriptional responses at single cell resolution *in vivo*

**DOI:** 10.64898/2025.12.23.696319

**Authors:** John A. Hawkins, Siamak Redhai, Svenja Leible, Mireia Osuna Lopez, Hilal Ozgur, Tianyu Wang, Michaela Holzem, Michael Boutros, Oliver Stegle

**Affiliations:** European Molecular Biology Laboratory (EMBL), Meyerhofstraße 1, 69117 Heidelberg, Germany; German Cancer Research Center (DKFZ), Division Signaling and Functional Genomics, Im Neuenheimer Feld 580, 69120 Heidelberg, Germany; German Cancer Research Center (DKFZ), Division Computational Genomics and Systems Genetics, Im Neuenheimer Feld 580, 69120 Heidelberg, Germany; Heidelberg University, Institute for Human Genetics, Department of Molecular Human Genetics & BioQuant, Medical Faculty Heidelberg, Im Neuenheimer Feld 267, 69120 Heidelberg, Germany

**Author notes:** Contributed equally (John A. Hawkins, Siamak Redhai).

**Keywords:** *In vivo* CRISPR detection, haplotyping, single-cell RNA sequencing, genotype-phenotype, computational genomics

## Abstract

CRISPR screens coupled with single-cell RNA sequencing are transforming high-throughput functional genomics. However, applications *in vivo* remain limited and are confounded by difficulties in identifying and genetically characterizing edited cells. Here we present scPT-seq, a single-cell RNA assay that resolves CRISPR-induced mutations at base-pair resolution and captures transcriptional responses in the same single cells *in vivo*. scPT-seq comes with a computational analysis suite enabling haplotype-resolved mutation detection and characterization of complex editing outcomes, including splice-junction variation. Applied to the *Drosophila* intestine, a highly regenerative tissue, scPT-seq distinguishes cell-autonomous from environmental effects by identifying mutant and wild-type cells within tissues, and reveals spatially organized compensatory mechanisms in response to mutations. By using editing outcomes as heritable clonal markers, we identified distinct intestinal stem cell populations with specialized differentiation trajectories. In summary, scPT-seq provides a versatile technology for dissecting gene function and lineage dynamics in complex tissues.

## Introduction

Understanding how genetic alterations give rise to distinct phenotypic traits lies at the heart of developmental biology, cancer research, and precision medicine. Genotype-phenotype relationships can be explored through population-scale studies that associate natural genetic variation with phenotypic diversity, or by experimentally introducing mutations^1^. CRISPR genome editing now enables systematic engineering of genetic variants at scale to determine how specific genetic changes influence cellular fitness and identity^2,3^. Combining CRISPR technology with single-cell RNA sequencing (scRNA-seq) has transformed this field, revealing how precise genetic changes reshape transcriptional programs and regulatory networks^4–8^.

Approaches for linking CRISPR perturbations to transcriptional phenotypes generally fall into two categories: detection of guide RNA expression and computational inference of perturbed state. Guide-based strategies have been utilized in large-scale pooled screens, but despite efficient delivery and expression of CRISPR components, a significant portion of sgRNAs fail to induce the intended edits, leaving the actual editing status unresolved^9,10^. Computational approaches can approximate perturbed states but remain indirect and sensitive to variability in editing efficiency^10–13^. In contrast, sequencing the edited locus establishes a direct connection between genotype and phenotype, revealing the full spectrum of editing outcomes and their cellular consequences. Recent methods that sequence DNA and RNA separately from the same cell therefore improve resolution but are limited in their ability to recover haplotypes or splicing changes and are often confined to a panel of transcriptional outputs^10,14,15^. Direct sequencing of edited RNA offers a promising alternative, as it builds on established, robust, and scalable whole transcriptome scRNA-seq workflows and can reveal posttranscriptional modifications such as splicing changes. Current approaches, however, have only profiled substitutional edits in an *in vitro* setting, leaving open how complex genetic mutations impact physiological systems^16,17^.

Here, we present Single-Cell Perturbation and Transcriptome sequencing (scPT-seq), an integrated experimental and computational framework that couples conventional high-throughput scRNA-seq with targeted long-read sequencing of the same cells to directly profile CRISPR-induced mutations and their transcriptional responses *in vivo*. scPT-seq yields the number, phasing, and base pair-level identity of edits in each cell, enabling differential expression analysis across subpopulations of cells grouped by genotypic status, markedly enhancing resolution and sensitivity. scPT-seq can distinguish genotypically-driven from environmentally-driven transcriptional changes by identifying genotypically wild-type cells within perturbed tissues. We applied scPT-seq to the *Drosophila* intestine, a highly regenerative organ, and uncovered spatially distinct regulatory programs that compensate for complex genetic edits. By using editing outcomes as clonal barcodes, scPT-seq permits lineage tracing and identifies distinct intestinal stem cell populations that generate region-specific cell types.

## Results

### scPT-seq enables direct detection of editing outcomes and transcriptome responses *in vivo*

scPT-seq extends established, robust, and scalable droplet-based single-cell RNA sequencing (scRNA-seq) workflows, such as 10X Chromium, to simultaneously capture whole-transcriptome profiles and editing outcomes from the same cells. In parallel to short-read sequencing of cellular mRNAs, targeted long-read sequencing on the cDNA pool enables direct recovery of the edited loci (Fig. 1a, Methods). Briefly, after cDNA synthesis, a nested PCR is performed using a universal 5′ primer (binding the Read1 adaptor) and a 3′ biotinylated adaptor downstream of the CRISPR target region (Fig. 1a, Methods). The resulting double-stranded amplicons contain the cell barcode, the UMI, and the edited region(s), and are enriched using a streptavidin pull-down. The enriched amplicons are further amplified using a nested primer and sequenced with long-read PacBio HiFi technology (Methods). The computational workflow of scPT-seq integrates genotypes with short-read expression data from the same cells, thereby providing for the first time a haplotype-resolved view of complex CRISPR-induced variants in RNA including insertions, deletions, substitutions, and splice-site alterations, linking these editing outcomes to genome-wide transcriptional responses and enabling clonal identification and lineage tracing (Supp. Fig. 1.1).

To maximize the information recovered from scPT-seq data, we devised a tailored computational workflow that is capable of reconstructing the full spectrum of genomic alterations (Supp. Fig. 1.2, Methods). First, pseudo-bulk aggregates of the long-read data are used to identify maternal and paternal segregating alleles for read phasing, and the long-read data from an unperturbed control sample is used to annotate all splice and missplice sites present in wild-type organisms (Supp. Fig. 1.3, Methods). Each long read is assigned to its corresponding cell barcode, phased using the nearest segregating variant, and scanned for mutations and novel splice events around the target locus (Supp. Fig. 1.2). At the cell–haplotype level, variants are classified as wild type, mutated, or uncalled based on coverage and allele-fraction thresholds, with only well-supported calls retained (Methods). A cell-haplotype is typically just a maternal or paternal chromosome and so we will typically use the term chromosome for simplicity, but note that in polyploid cells such as enterocytes each chromosome may have multiple physical copies. Importantly, the haplotype-resolution allows scPT-seq to identify missing haplotypes, providing a unified, guide-independent view of editing and splicing changes from the same molecule.

**Figure 1:**
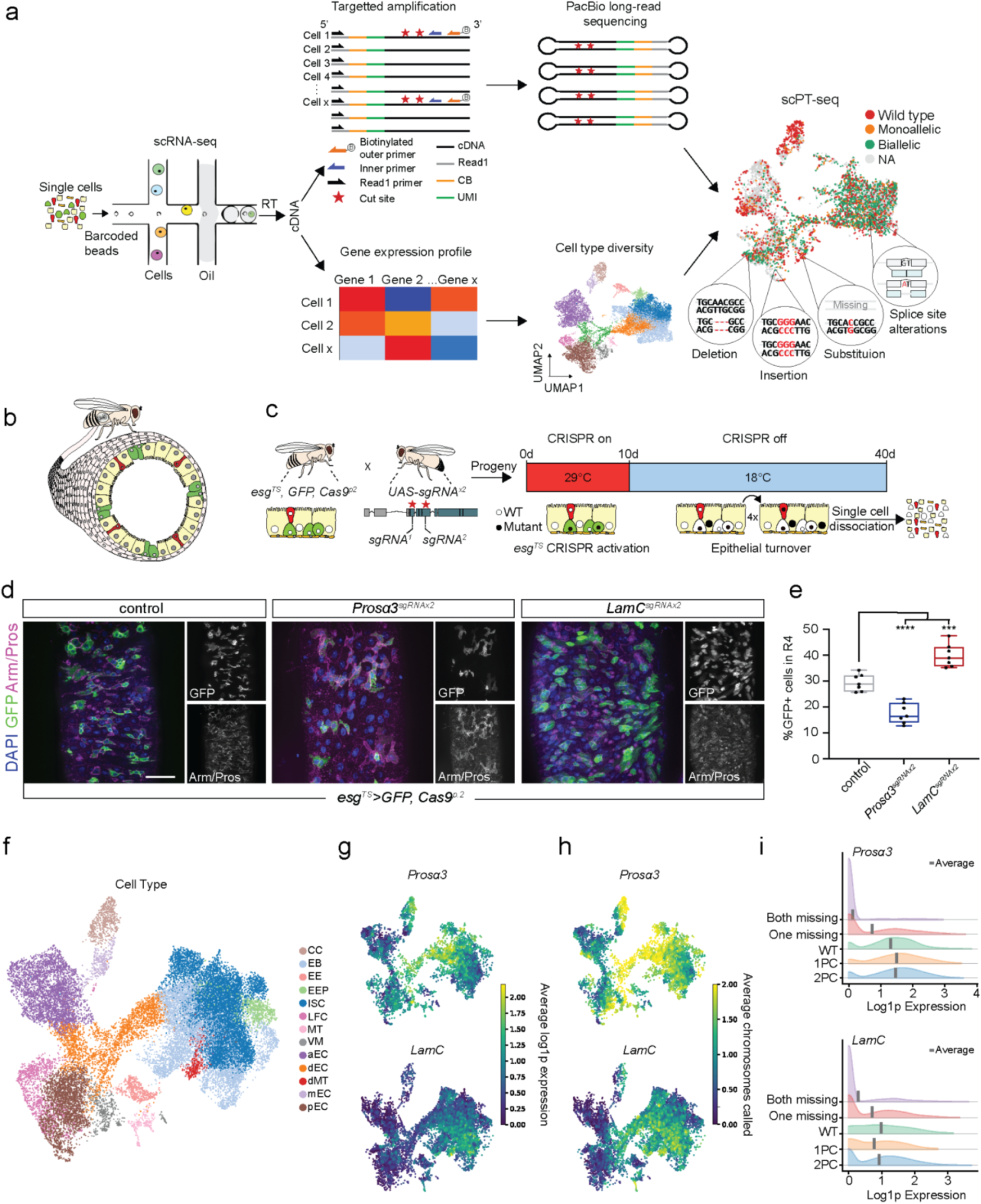
Direct detection of editing outcomes and transcriptome responses using scPT-seq. **a.** scPT-seq experimental overview. Single cells are processed with droplet-based scRNA-seq. The resulting full length cDNA library is split into two aliquots. The first is fragmented and processed with short-read sequencing to obtain whole transcriptome expression data in individual cells. For the second aliquot, the target site of interest is enriched with biotinylated primers and sequenced using long-read PacBio HiFi technology, ensuring cell barcodes and genomic edits are captured on the same molecules. This pipeline identifies both mutations and whole transcriptome data within the same single cell. **b.** The Drosophila midgut contains a variety of cell types with function-dependent spatial organization. Green: ISC, yellow: EC, red, EE, white: surrounding cells, grey: nuclei. **c.** Genetic crossing scheme for CRISPR activation. Transgenic female flies carrying the esg^TS^ expression system, enabling temperature-inducible activation of GFP and Cas9 in progenitor cells, are crossed with male flies carrying the two sgRNAs for the gene of interest. After progeny have reached adulthood with normal development, they were moved to 29*°*C for 10 days to induce expression of sgRNAs, and GFP in progenitor cells. The flies were then moved back to 18*°*C for 30 days to reach steady state. During this time, the epithelium turns over approximately four times, enabling progenitor-derived mutations to spread throughout epithelial cells. The midgut is disassociated into single cells and processed with scPT-seq. **d.** Confocal images of the midgut demonstrating that CRISPR mutagenesis to *Prosα3* reduces GFP+ progenitor cell number, whereas LamC perturbation increases the progenitor population. For imaging, flies were moved to 29*°*C for one day following the 30 day steady state period to reactivate GFP in progenitor cells. Green: GFP, blue: DAPI, magenta: Arm/Pros. Scale bar 100 μm. **e.** Quantification of the progenitor population in (d). **f.** UMAP of all cells in all samples, colored by cell type. CC: copper cell, EB: enteroblast, EE: enterocyte, EEP: EE progenitor, ISC: intestinal stem cell, LFC: large flat cells, MT: malpighian tubule, VM: visceral muscle, aEC: anterior EC, dEC: differentiating EC, dMT: differentiating MT, mEC: middle EC, pEC: posterior EC. **g.** Average log1p expression of the gene of interest averaged over a neighborhood of 10 cells in its corresponding samples. **h.** The average number of chromosomes per cell with mutations identified from long read sequencing, averaged in a neighborhood of 10 cells around each cell. All cells, including polyploid cell types, are considered to have two chromosomes here, corresponding to the two chromosomes (and their corresponding mutations) in the parent stem cells. **i.** log1p expression distributions (short read sequencing) stratified by the mutation status (long read sequencing). Each cell is considered to have two chromosomes, similar to (g), of which one or both can be missing. Cells with mutation status identified in both chromosomes are divided into wild-type (WT), one perturbed chromosome (1PC), and two perturbed chromosomes (2PC). The average log1p expression for each category is shown as a vertical grey bar.

To demonstrate the power of scPT-seq in a complex and dynamic tissue, we applied it to the *Drosophila melanogaster* midgut, a well-established model for studying tissue homeostasis and regeneration^18–20^. The midgut is a self-renewing epithelium directly exposed to the external environment and analogous to the mammalian intestine, with well-defined cellular diversity, lineage hierarchies and spatial organization that are essential for tissue function (Fig. 1b). These core features make the midgut a representative and tractable system for testing how scPT-seq captures cellular states, lineage relationships, and spatially organized programs in a complex tissue. scPT-seq is particularly well suited for this system because conventional guide-capture perturbation readouts do not apply owing to the timeframe needed to produce phenotypes^21^. Captured midgut cells include cells along the full lineage hierarchy: intestinal stem cells (ISCs)^22,23^ drive tissue turnover by giving rise to enteroblasts (EBs) and enteroendocrine precursors (EEPs), collectively termed progenitors, which subsequently differentiate into subclasses of enterocytes (ECs) and enteroendocrine cells (EEs), respectively (Fig. 1b).

Using a temperature-inducible, progenitor-specific driver, we are able to control the time and cell type for CRISPR mutagensis^21,24^. We restricted CRISPR mutations to the adult stage, ensuring that editing occurred only in the progenitor cell population and that transcriptional readouts reflected post-developmental effects. Specifically, we induced *sgRNA^x^*^2^ and CRISPR-Cas9 expression for ten days at a permissive temperature and subsequently allowed the midgut to undergo approximately four rounds of turnover at restrictive temperature before utilizing scPT-seq (Fig. 1c). We perturbed two genes with opposing effects on progenitor dynamics: *Prosα3*, a regulatory α-subunit of the 20S proteasome that reduced progenitor numbers when perturbed, and *LamC*, a nuclear intermediate filament whose mutagenesis led to an expansion of the progenitor population (Fig.1d,e). Both are important for epithelial homeostasis and are frequently dysregulated in human cancers, making them suitable candidates for scPT-seq^25,26^. Notably, despite the use of nominally inbred stocks, heterozygosity, estimated at roughly one variant per 400 bp in the exome in our data, provided sufficient sequence diversity for haplotype phasing of long-read data. This enabled allele-specific analysis of editing outcomes (Methods).

We captured a total of 26,068 cells across the two perturbations, *Prosα3^sgRNAx^*^2^ and *LamC^sgRNAx^*^2^, including two biological replicates per perturbation and two independent control experiments without perturbation, with an average of 325 long reads per cell covering the edited locus and encompassing all major intestinal cell types (Fig. 1f and Supp. Fig. 1.3). Both *Prosα3* and *LamC* displayed pronounced cell-type–specific expression patterns (Fig. 1f,g). We achieved robust mutation detection across cell types, exceeding the sensitivity of short-read scRNA-seq (Fig. 1g-i). We detected at least one chromosome in 77% of cells in *Prosα3* and and 49% in *LamC*, including many cells with zero measured expression from short reads (Fig. 1i, Supp. Fig. 1.4). Of note, both perturbations are in the first exons of multi-exon transcripts subject to nonsense mediated decay, yet cells with perturbed *LamC* did not show substantially decreased expression, and observed expression actually increased for perturbed *Prosα3* (Fig. 1i). Collectively, these results confirm that scPT-seq achieves high sensitivity across diverse cell types, enabling reliable detection of edited transcripts.

### scPT-seq reveals previously masked diversity of editing outcomes

First, to understand the spectrum of edits that occur, we aggregated detected chromosomes for *Prosα3* and *LamC* perturbed cells respectively (Fig. 2a), finding that the variant frequency was markedly concordant across biological replicates (Fig. 2b, Supp. Fig. 2.1). The most common edits in both *Prosα3* and *LamC* were short deletions, though in each case 3 of the top 10 edits were splicing related, either intron retention from complete loss of splicing or changes to splice junction locations (Fig. 2a). For *Prosα3*, aside from intron retention, the three most common edits were in-frame: a 12 bp deletion, a 3 bp deletion, and an 18 bp splicing junction site shift. These edits collectively account for 34% of the total alterations, whereas the mutation spectrum of *LamC* was more uniformly distributed, with no particular edits present in such a large fraction of chromosomes (Fig. 2a). The most common alteration for *LamC* and second most common for *Prosα3* was intron retention (Fig. 2a), which can result from any mutation that disrupts the splice site. The prevalence of splicing changes for both targets exemplifies how scPT-seq captures post transcriptional modifications induced by CRISPR edits, which cannot be captured by DNA-centric approaches (Fig. 2c, Supp. Fig. 1.1). These insights into the mutational spectrum highlight the power of scPT-seq to resolve editing diversity *in vivo*.

We next quantified editing zygosity across all intestinal cell types. Across cell types, *Prosα3* perturbation produced mixed zygosity profiles, with the highest editing frequencies in progenitors, EEPs, EEs, and posterior enterocytes (pECs), whereas anterior enterocytes (aECs), middle midgut enterocytes (mECs), and copper cells (CCs) remained largely wild type (Fig. 2d). In contrast, *LamC* perturbation yielded predominantly biallelic edits across nearly all populations, with reduced editing limited to mECs and CCs (Fig. 2e). Together, these results reveal cell type– and region–specific variation in editing efficiency, highlighting the influence of tissue context on CRISPR mutagenesis.

**Figure 2:**
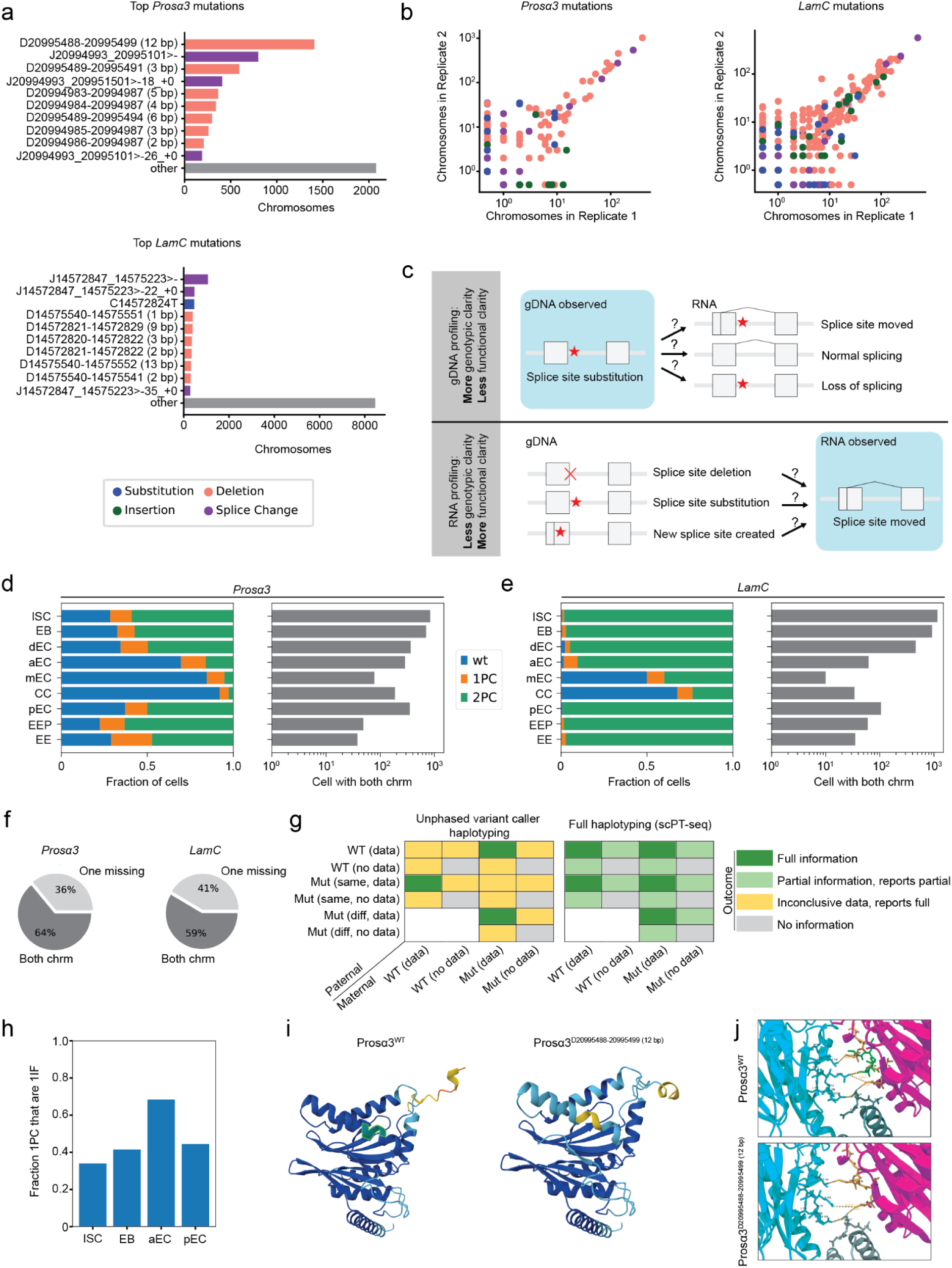
scPT-seq reveals previously masked diversity of editing outcomes. **a.** The top 10 mutations from each perturbation, and their total chromosome count. Colors give the type of mutation, shown in the legend below. Ja-b>-indicates intron retention (loss of the indicated splice junction). Full abbreviation specifications in Methods. **b.** Reproducibility of mutation spectrum. For each mutation observed, the number of chromosomes with that mutation is plotted for each of the two biological replicates. Colors same as (b). **c.** Examples of gDNA- and RNA-based profiling strengths and limitations. gDNA profiling identifies the genotype directly, but does not clearly identify the consequences for splicing. RNA profiling may miss genotypic mutations removed via splicing, but directly identifies splicing changes and the resulting functional RNA sequence. **d.** Cell mutation status and cell count statistics by cell type for *Prosα3*. Left panel: The fraction of cells with mutation status identified on both chromosomes whose cell mutation status is wt, one perturbed chromosome (1PC), or two perturbed chromosomes (2PC), stratified by cell type. Right panel: The number of cells per cell type with mutation status on both chromosomes identified. **e.** Same as (d) for *LamC*. **f.** Pie charts showing fraction of cells with mutation calls with data from one or both chromosomes. **g.** Comparison of haplotyping output for cells with data from one or both chromosomes using unphased variant caller-based haplotyping or full haplotyping. Ground truth of mutation status and sequencing sampling status for maternal and paternal chromosomes are given on the x- and y-axes. Mutation statuses: WT, Mut (same) for same mutations on paternal and maternal, and Mut (diff) for different mutations on the two chromosomes. Sequencing sampling statuses: data or no data. **h.** For each of the four most frequent cell types in our data, the fraction of cells with one perturbed chromosome (1PC) where that chromosome mutation is in-frame (1IF). **i.** Structural AlphaFold predictions of wildtype *Prosα3* and the variant containing the most common 12 bp deletion. Both structures are colored according to the pLDDT score, reflecting prediction confidence. The helix of interest shows a confidence value >50 in both models. In the wildtype structure, the nucleotides affected by the mutation are highlighted in green (DAN**VLTS**ELR). **j.** Predicted interactions between *Prosα3* (magenta), *Prosα*2 (blue) and *Prosβ3* (dark green). Potential interaction sites were manually selected and displayed in stick representation. The mutated amino acids VLTS are shown in green. Following mutation, distances between potential interaction sites increase in all illustrated cases, while interactions involving the mutated amino acids are lost. Both structures exhibit only minor positional changes, suggesting that functional interaction is still likely.

A unique advantage of scPT-seq is its ability to disambiguate missing alleles (Fig 2f,g). Previous methods either do not haplotype or use unphased variant caller-based haplotyping^10,14,15,17^, which can be a significant source of error. WT cells and cells with the same edit on both chromosomes are indistinguishable from cells with missing chromosomes using unphased variant caller-based methods (Fig 2g). This distinction is critical *in vivo*, where incomplete capture or stochastic allele loss is frequent. scPT-seq explicitly labels such cases as “missing,” thereby preventing such errors. We find this is a significant effect, where 36% of *Prosα3* and 41% of *LamC* cells with identified edits in our data would be subject to this source of error in unphased variant caller-based methods (Fig. 2f).

The overall patterns we observed in the identities and distributions of edits hinted at another important possibility: selection of different mutations and in different cell types during the 30 days of cell turnover following CRISPR perturbation. For example, the difference between the zygosity profiles of aECs between the *Prosα3* and *LamC* samples is striking (Fig 2d,e). mEC and CC cells are broadly WT in both perturbations, which could possibly be due to a known tendency toward senescence of progenitor cells in this region, or some other tissue specific mechanism preventing perturbation generally^27^. But the fact that aECs are almost completely biallelically mutated in *LamC* means that the CRISPR machinery and cell turnover works efficiently in this cell type, so the difference in sgRNA target sequence alone has caused a systematic reduction in perturbations in the *Prosα3* sample in a cell type- and region-specific manner. We followed up on this observation by looking at the kinds of mutations in aECs with edits. Restricting to monoallelically edited cells to reduce noise, we looked at the fraction of cells where the *Prosα3* perturbation was in-frame across our four most common cell types. We found that aECs are highly enriched for in-frame mutations (Fig. 2h), consistent with cell type- and region-specific selection for functional or partially functional *Prosα3*. We did not, however, find elevated signatures of apoptosis in aECs by mutation status (Supp. Fig. 2.2), suggesting the perturbations could be causing a region-specific change in the Waddington landscape rather than cell type-specific increase in perturbation lethality^28^.

As a second example of potential selection, we examined the most frequent edit in the Prosα3 sample: an in-frame 12 bp deletion. This mutation removes four amino acids from an α-helix (residues 80–101), deleting one full α-turn (Fig. 2i). AlphaFold2 Multimer modeling of the 20S proteasome with and without the deletion showed loss of the Q112 (Prosα2)–S88 (Prosα3) interaction and altered distances among local interacting residues (Fig. 2j). Despite these local perturbations, the overall quaternary structure remained intact, suggesting the mutation could affect local flexibility rather than core assembly (Supp. Fig. 2.3). Such subtle changes may be tolerated while modulating proteasome activity. Together, these results illustrate how scPT-seq links fine-scale mutational patterns to structural and functional hypotheses.

### scPT-seq increases resolution for differential expression analysis and uncovers cell-type specific dosage response

By recovering the full perturbation status of each cell, scPT-seq enables more granular comparisons than previous methods without direct perturbation sequencing. *In vivo* methods in particular often compare perturbed samples against independent control samples without any perturbation information at the single cell level. gRNA capture methods, meanwhile, report only a simple yes/no measurement of whether a cell is perturbed without chromosome-level information, and furthermore rely on two noisy assumptions: observed gRNA means a cell was edited and lack of observed gRNA means a cell was not edited. The noise in these assumptions requires larger numbers of cells to find meaningful signals, and in cases of low editing efficiency the signal can entirely be obscured for biological reasons. In contrast scPT-seq provides full, chromosome-level, base pair-resolution detail of the mutation status of each cell. Consider the first simple binning of these results into wild type, monoallelic (1PC), and biallelic (2PC) perturbations. This already combinatorially expands the number of possible subsample comparisons to seven (Fig. 3a) and enables dosage response analysis.

First, considering the *Prosα3^sgRNA×^*^2^ perturbation, we systematically considered all possible comparisons of the control sample and the perturbed sample stratified into WT, 1PC, and 2PC cells, conducting 54 differential expression (DE) comparisons across 11 cell types, and examined 1,431 pairwise correlations of DE results (Fig 3b). Broad patterns emerge across comparisons. Diagonal lines in the correlation matrix show that different comparisons performed in the same cell type tend to correlate, and ISC and EB progenitor cells also tend to have elevated correlation, for example.

**Figure 3:**
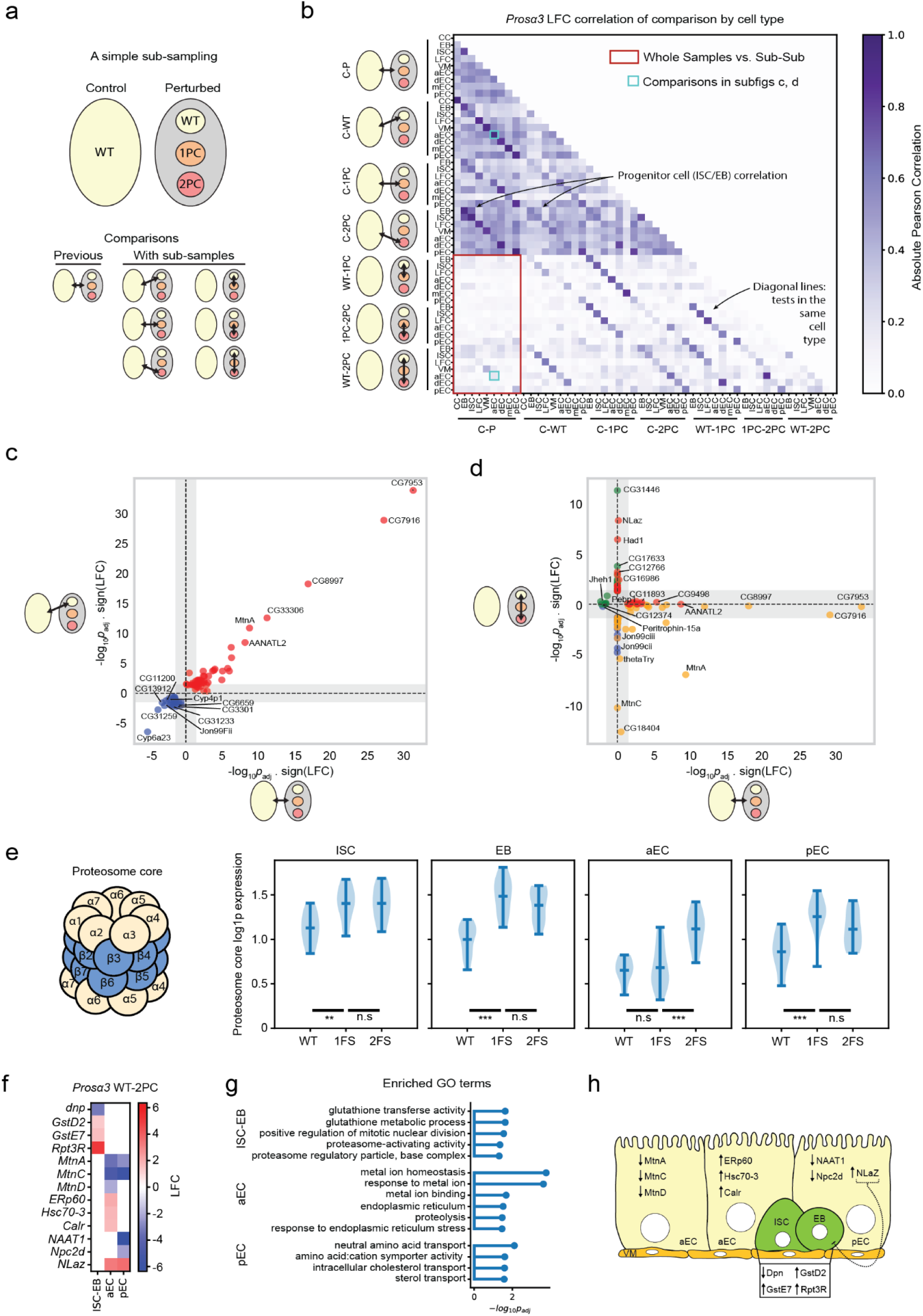
Novel comparison opportunities for differential expression analysis and previously hidden effects. **a.** A simple subclassification strategy for the perturbed cell statuses from scPT-seq and the several new differential expression (DE) comparisons made possible. 1PC: one perturbed chromosome, 2PC: two perturbed chromosomes. **b.** Correlation plot for all comparisons shown in (a), stratified first by comparison strategy and then by cell type. Red bounding box: correlations between the whole-sample comparison and comparisons within subsamples of perturbed cells. **c.** Volcano vs. Volcano plot showing C-WT vs. C-P comparison in aEC cells. Grey bars indicate non-significant results along respective axes. p_adj_: BH FDR corrected p-values, LFC: log fold change. Red: up-regulated in both comparisons, blue: down-regulated in both comparisons, green: up on y- and down on x-axis, yellow: down on y- and up on x-axis. **d.** Same as (c) but for WT-P2C vs. C-P. **e.** Dosage effects of the proteasome core across the major cell types. 1FS: one wt chromosome and one with frame shift mutation, 2FS: frame shift edits on both chromosomes. ** and ***: p-value less than 0.01 and 0.005 respectively. n.s. not significant. **f.** Expression log fold change for genes of notable biological interest in our major cell types. **g.** Selected enriched GO terms for results in (f). **h.** A model of genes of interest affecting stem cell proliferation.

The most important patterns in our comprehensive profiling, however, reveal that scPT-seq avoids confounding of differential expression analysis by using true internal controls rather than a matched control sample, or even guide capture-identified controls. All subsets of cells in the perturbed sample of any genotype show highly correlated transcriptional changes when compared with the control sample (Fig. 3b, darker upper-left triangle). For example, this means the results are very similar whether we compare WT versus control or the whole perturbed sample versus control (Fig. 3c). This is true to a lesser degree even if the comparisons are in different cell types (Fig. 3b). Upregulation of genes such as *GstD2* and *MtnA*, along with several uncharacterized transcripts, reflect batch-associated and general stress responses rather than mutation-specific effects. In contrast, comparisons between different genotypes within the perturbed sample, such as 2PC-edited versus WT cells, produce almost entirely orthogonal DE signatures with the perturbed sample-versus-control analysis (Fig. 3d). This effect is particularly striking in aECs, in which a high percentage of perturbed sample cells are WT (Fig. 2d), but the loss of correlation between comparisons is true across cell types (Fig. 3b, red outline). The 2PC versus WT cells comparison identifies effects expected from the perturbation itself, including compensatory proteasome core upregulation and activation of secreted growth inhibitors like *NLaz*^29^, while none of these key responses are apparent in the bulk perturbed-versus-control analysis, which is instead dominated by nonspecific stress-related genes (Fig. 3d). Taken together, we find that comparisons against the control sample here return many significant genes, but obscure the biological results we originally set out to find, and can thus mislead interpretation without mutation status explicitly incorporated. The WT versus control comparison may be interesting in its own right to study environmental effects, for example, but knowing the mutation status is crucial for that interpretation as well.

Furthermore, guide capture methods would not be expected to address the signal obscuration problem here. In the *LamC^sgRNA×2^* experiments nearly all aEC cells have biallelic edits, so the CRISPR editing machinery is effective in this cell type (Fig. 2e). The only difference in experimental setup is the sgRNA target sequence, meaning scPT-seq here identifies majority WT genotypes in cells where gRNA capture would be expected to report nearly fully perturbed cells.

scPT-seq also enables quantitative analysis of dosage-dependent transcriptional regulation. To explore this, we focused on the 20S proteasome complex to probe compensatory mechanisms upon *Prosα3* disruption. At the cell type level, distinct transcriptional patterns emerged, revealing context-dependent modes of compensation (Fig. 3e). In ISCs and EBs, *Prosα3* disruption with frameshift mutations elicited a marked upregulation of proteasome core genes, with significant increases from fully WT to one frameshift-mutated chromosome and one WT chromosome (1FS) that plateaued with frameshift mutations in both chromosomes (2FS), indicating that progenitor populations mount a similar compensatory response to reduced gene dosage. Notably, such compensatory activation of proteasome subunits has been reported previously following disruption of proteostatic checkpoints^25^, underscoring that scPT-seq captures biologically meaningful transcriptional signatures. In contrast, differentiated enterocytes exhibited opposing trends: aECs required biallelic *Prosα3* loss (2FS) to trigger proteasome core gene upregulation, whereas pECs responded already at the monoallelic state (1FS) (Fig. 3e). Together, these results demonstrate that dosage-dependent transcriptional compensation is not uniform but instead spatially distinct and cell type–specific within the midgut epithelium.

Next, we leveraged large language models to systematically look for patterns across the many differential expression comparisons across all midgut cell types to look for gene signatures associated with our perturbations. To do this, we first generated a large report which contained, for all comparisons in all cell types: DE genes and effect sizes, enriched GO terms, and plain text descriptions of the function of each DE gene from FlyBase^30^ (Methods). We then confirmed the results of the LLM analysis by manual inspection. In *Prosα3*, this analysis revealed an enrichment of detoxification and stress-response genes within biallelically edited ISC/EBs, accompanied by downregulation of self-renewal regulators, including the transcription factor *deadpan*, which is known to be required for progenitor proliferation in the midgut^31^ (Fig.3 f,g). In pECs, *Prosα3* loss resulted in reduced expression of genes involved in amino acid and sterol transport, consistent with impaired absorptive function, and increased expression of the secreted insulin-signaling antagonist *NLaz*, suggesting potential non-autonomous suppression of progenitor proliferation through reduced insulin pathway activity (Fig. 3f,g). In aECs, biallelic edits in *Prosα3* caused strong downregulation of genes involved in metal sensing, binding, import, and storage indicating decreased micronutrient sensing. Concurrently, we observed upregulation of numerous endoplasmic reticulum stress–responsive genes, likely evoked due to impaired proteasomal function (Fig. 3f,g). Across cell types, we also detected enrichment of terms related to proteasome-activating activity and proteolysis, consistent with a compensatory transcriptional response to *Prosα3* disruption (Fig. 3f,g).

For *LamC^sgRNAx2^*, since edits were predominantly biallelic, we directly compared biallelically edited cells with wild-type controls. This revealed cell type-specific responses to *LamC* perturbation (Supp. Fig. 3.1). In progenitor cells, we observed a coordinated upregulation of Notch pathway targets, including *E(spl)m7-HLH*, *E(spl)m8-HLH*, and *E(spl)m6-BFM*, and a decrease in the expression of the transcription factor *lozenge* (*lz*), which has been previously reported to trigger non-autonomous ISC proliferation^32^. Moreover, the Toll signaling transcription factor *Dif*, genes governing cytoskeletal organization and cell adhesion, and detoxification-associated genes were downregulated, indicating a broad suppression of immune responsiveness accompanied by impaired cellular structural integrity and reduced capacity to mitigate physiological stressors. aECs responded to *LamC* perturbation by downregulating digestive enzymes (Lysozymes) and immune-related genes, as well as transporter proteins (*dmGlut* and *LpR1*). Like progenitor cells, both aECs and pECs upregulated Notch signaling related genes, indicating that *LamC* perturbation similarly impacts this pathway across different cell types. Additionally, aECs increased expression of the JAK–STAT ligand *Upd2*, previously linked to non-autonomous stimulation of progenitor proliferation under stress conditions^33^. Similarly, pECs upregulated *Upd3* and *Rhomboid*, the latter required for EGF ligand secretion and subsequent progenitor proliferation^34,35^. These findings suggest that the excessive proliferation triggered by *LamC* perturbation in progenitors is, in part, driven by non-autonomous JAK–STAT and EGF signaling.

Together, these results demonstrate that scPT-seq enables high-resolution mapping of cell type–specific transcriptional consequences of perturbation, providing a mechanistic framework for the observed changes in the progenitor cell population following *Prosα3* and *LamC* mutagenesis (Fig. 3h, Supp. Fig. 3.1).

### scPT-seq enables clonal analysis and spatially informed lineage tracing and DE analysis

Perturbation-induced changes in progenitor cell proliferation is predominantly studied in the posterior midgut, but spatial differences along the length of the gut are of growing interest in the community^36^. The *Drosophila* midgut comprises a variety of differentiated cell types with distinct molecular and functional identities that are spatially organized along the anterior–posterior axis (Fig. 4a), yet accurately resolving the position of progenitor cells using scRNA-seq remains a challenge due to their transcriptional similarity (Fig. 4b).

To study the spatial differences of progenitor cells, we leveraged scPT-seq to trace lineages and transfer spatial information between cell types. Our experimental design restricts expression of the CRISPR perturbation machinery to the progenitor cell population and produces a final population of edited cells of each cell type, including progenitor cells, through descent during approximately four full rounds of cell turnover (Fig. 1c). Each unique combination of edits thus serves as a heritable clonal marker, enabling reconstruction of lineage relationships directly from the editing outcomes. This requires each combination of edits is in fact unique, so we implemented a data-driven model to assess uniqueness and filtered out combinations of edits with high probability of occurring in our dataset by two or more original editing events (Methods). After filtering, we labeled sets of cells with the same unique combination of mutations as clonal cell populations. Each clone is in a particular location in the tissue, which thus allows us to transmit spatial information between cells in the same clone (Fig. 4b).

**Figure 4:**
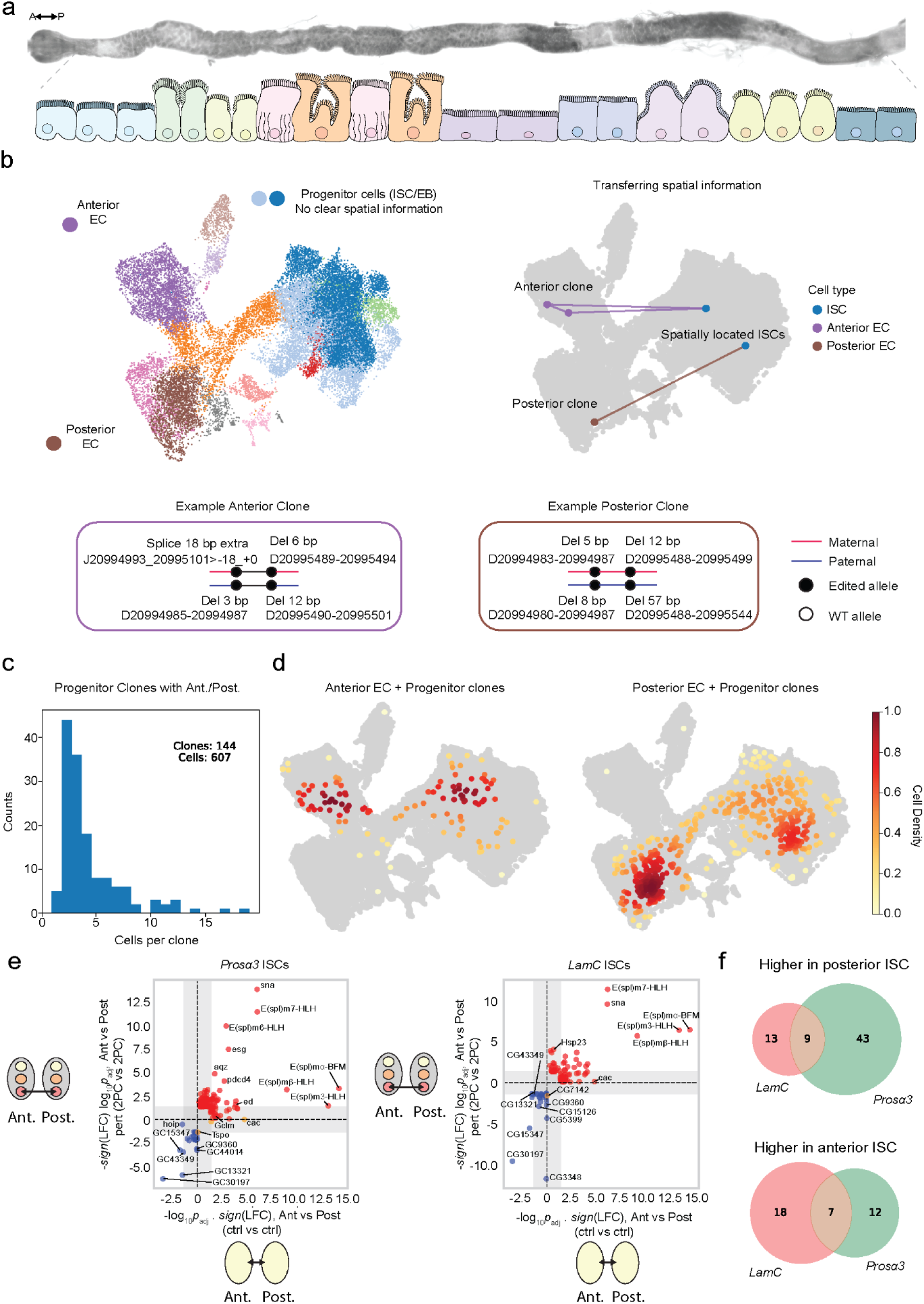
Clonal analysis, spatially informed lineage tracing, and spatial DE analysis. **a.** Cell types and their spatial position along the anterior-posterior axis of the midgut. **b.** Spatially locating progenitor cells via clonal analysis. Progenitor cells lack clear spatial information from their expression profiles alone, in contrast with ECs. The mutations present in each cell serve as a marker identifying clones of cells descended from one original edited stem cell. We identify clones of cells that contain both progenitor cells and either anterior ECs or posterior ECs, and thereby transfer spatial information from ECs to progenitor cells. **c.** Histogram of cells per clone for clones that contain progenitor cells and either aECs or pECs. **d.** All clones in (c), plotted on UMAPs for clones with anterior ECs or posterior ECs respectively. Color: cell density. **e.** Volcano vs. Volcano plots showing anterior-posterior DE analysis in 2PC perturbed cells vs. the same in control samples. **f.** Venn diagrams of DE results unique to perturbed comparison.

The high throughput nature of scPT-seq allows us to find many small clonal populations of cells and aggregate their spatial information to study larger scale effects. Using the method described above, we identified clones of cells containing both our progenitor cells of interest and either anterior ECs (aECs) or posterior ECs (pECs) for spatial registration (Fig. 4b). In our experimental setup, we capture on average only about 1 in 300 cells from each original midgut, yet we were still able to identify 144 clones with this particular spatial link, comprising a total of 607 cells (Fig. 4c). Most clones contained fewer than five cells. The combination of all these clones allowed us to see global lineage patterns of anterior and posterior ISCs that differentiate into region-specific ECs in the midgut. In particular, we identified subclusters of the data enriched for spatially registered anterior and posterior ISCs. (Fig. 4d and Supp. Fig. 4.1). With these subclusters identified, we could look for spatial ISC transcriptomic expression patterns. Our results recovered known differences between anterior and posterior ISCs, such as enrichment for genes belonging to the Notch pathway and higher *esg* expression in posterior cells^37^ (Fig. 4e). Our findings were consistent between both the *Prosα3* and *LamC* datasets, demonstrating distinct regional identities for ISCs. These results demonstrate that scPT-seq can recover spatial information for transcriptionally similar cell types, enabling integrated analysis of genotype, expression, lineage, and spatial context from a single experiment.

We next asked how ISCs from distinct midgut regions respond to genetic perturbation by identifying region-specific differentially expressed genes following *Prosα3* or *LamC* mutagenesis. Under *Prosα3* perturbation, 52 genes were significantly upregulated in posterior relative to anterior ISCs (Fig. 4e,f), indicating region-dependent molecular responses to proteasome dysfunction along the gut axis. We observed strong overrepresentation of transcriptional and chromatin regulators (e.g., *bun*, *cnc*, *Tet*, *lola*, *Blimp*-*1*, *sd*) as well as genes involved in signal transduction and developmental patterning (*N*, *Egfr*, *sgg*, *brk*, *REPTOR*) in posterior ISCs. Consistent with these patterns, GO terms included stem cell homeostasis, negative regulation of transcription, and multiple signalling pathways. In contrast, anterior ISCs upregulated 19 genes relative to posterior ISCs in response to *Prosα3* editing, with GO terms associated with actin cytoskeleton and cell-junction organization, mitochondrial electron transport, and DNA repair. Region-specific gene-expression patterns between anterior and posterior ISCs were also observed following *LamC* perturbation and showed partial overlap with the *Prosα3* response (Fig. 4e,f). For example, ∼40% of genes upregulated in *LamC^sgRNAx2^* posterior ISCs were shared with *Prosα3* perturbation. However, *LamC^sgRNAx2^* also induced distinct transcriptional programs, including increased expression of stress-response and metabolic genes. In anterior ISCs, *LamC* perturbation led to enrichment of genes related to ribosome biogenesis, mitochondrial function, membrane transport, and apoptosis. Taken together, these findings show that scPT-seq not only recovers position information of ISCs but also uncovers the region-dependent molecular programs that emerge under perturbed conditions.

## Discussion

Determining how precise genomic edits alter cell states requires methods that can identify the exact mutation in each cell, link that genotype to a comprehensive molecular phenotype, and operate within complex tissues where cell identities and spatial context are critical for homeostasis. Here, we introduce scPT-seq, an approach that fulfills these criteria and further expands them by haplotyping mutations, identifying changes in splice junction sites and enables spatially informed analysis of genome editing outcomes at single-cell resolution. Applied *in vivo*, scPT-seq identified and differentiated between transcriptional programs driven by genotypic changes as opposed to those arising from the environment by identifying genotypically wild-type cells within perturbed tissues. By incorporating the number, phase, and type of edits in each cell, scPT-seq found spatially organized dosage response compensatory mechanisms across the midgut. And by leveraging editing outcomes as heritable clonal markers, scPT-seq revealed lineage relationships, spatial identity, and region-dependent perturbation responses among transcriptionally similar cell types.

Pooled CRISPR–scRNA-seq screens have transformed functional genomics, but most implementations still infer which cells are edited indirectly, through sgRNA detection, computational prediction, or reporter constructs, rather than by sequencing the edited locus itself^3–5^. This indirect inference limits resolution when editing efficiency is variable, heterogeneous, or yields multiple alleles from a single guide. Recent innovations have begun to bridge this gap from several directions. Combining short and long-read sequencing demonstrated that CRISPR edits can be recovered for highly expressed genes and that one could identify simple CRISPR base-editor mutations in cell lines^16,17^. Complementary joint DNA–RNA strategies such as SDR-seq and CRAFTseq directly capture genomic edits and associate them with transcriptomic and, for CRAFTseq, proteomic phenotypes^10,14^. However, these methods are limited in their throughput of cells or genes, their ability to resolve complex alterations such as splicing changes, their ability to fully haplotype mutations, and have so far only been applied *in vitro*. scPT-seq enables per-cell, haplotype-resolved genotyping of complex editing outcomes, including mixtures of small/large indels, substitutions and splice-junction changes, by integrating targeted long-read amplicons with droplet-based transcriptional profiling of thousands of cells. By enriching edited loci through a biotin-streptavidin pull-down, scPT-seq captures targets with varying expression level and has the capacity to be multiplexed.

Applied to the *Drosophila* midgut, scPT-seq uncovered previously masked heterogeneity of editing outcomes: a large variety of edits per guide, reproducible across replicates, and with clear cell-type specific distributions. By phasing edited reads, we disambiguated true monoallelic edits from allelic dropout/missing chromosomes, a major potential source of noise in single-cell unphased variant calling. We further looked at predicted structural alterations of a prevalent *Prosα3* mutation, uncovering potential selective pressures that compensate for perturbing an essential gene. Additionally, we uncover how genomic edits drive distinct transcriptional programs and reshape epithelial niche feedback signals to impact the intestinal progenitor population.

Although scPT-seq offers substantial advantages, several technical considerations remain that present opportunities for further refinement. scPT-seq currently depends on the presence of heterozygous variants or local sequence diversity to achieve haplotyping, which may limit its application in fully inbred systems unless complemented by synthetic SNP or barcodes. In addition, long-read sequencing remains costlier and lower throughput than short-read platforms, though the targeted enrichment strategy used in scPT-seq mitigates this limitation by focusing depth on informative molecules and providing complete coverage of edited sites from relatively few reads. As sequencing technologies continue to improve in cost, accuracy and read length, the scalability of scPT-seq will further increase. Finally, while scPT-seq directly links genotypes to transcriptional profiles, it does not yet capture other molecular modalities, such as chromatin accessibility and protein expression, that would enable a more comprehensive understanding of how specific edits reshape cellular states within their native tissue context.

## Methods

### Fly husbandry

*Drosophila* stocks were maintained on a 12:12 hour light–dark cycle at standard laboratory conditions. Flies were raised on standard cornmeal-based food, prepared per liter with the following ingredients: 44 g syrup, 80 g malt, 80 g corn flour, 10 g soy flour, 18 g yeast, 8 g agar, 2.4 g methyl-4-hydroxybenzoate, 6.6 mL propionic acid, and 0.66 mL phosphoric acid. For experiments involving temperature-sensitive transgene induction using *Gal80^TS^*, parental flies were reared at 18 °C, and progeny were shifted to 29 °C after eclosion to activate transgene expression. For CRISPR-Cas9 mutagenesis, flies were first shifted to 29 °C after eclosion for 10 days then shifted to 29 °C for a further 30 days. In all experiments, mated female flies were used and transferred to fresh food every two days.

### Fly stocks

The following fly lines were used in this study: *esg^TS^, UAS-GFP* (gift from B. Edgar), *esg^TS^, UAS-GFP, UAS-Cas9^p2^* (Boutros Lab), *UAS-Prosalpha3^sgRNAx2^* (VDRC: v341974), *UAS-LamC^gRNAx2^* (VDRC: v341947).

### CRISPR-Cas9 experiments

For progenitor-specific mutagenesis with *Cas9^p2^*, sgRNA lines were selected with close proximity to the 5’ end of targeted genes. Newly eclosed flies were shifted to 29°C for 10 days then 18°C for a further 30 days, as previously described (Bahuguna et al, 2021).

### Dissection and immunohistochemistry

Adult female flies were shifted to 29 °C for 1 day to induce progenitor-specific GFP expression and starved for 3 hours prior to dissection to minimize intestinal luminal content. Intestines were dissected in PBS (Phosphate Buffered Saline; Sigma-Aldrich, P3812-10PAK) and transferred to polylysine-coated slides. Tissues were fixed in 4% paraformaldehyde (diluted from 16% stock; Thermo Scientific) in PBS for 60 minutes. Following fixation, samples were washed in PBST (PBS with 0.1% Triton X-100) for 30 minutes, then blocked in PBSTB (PBST with 1% Bovine Serum Albumin) for 30 minutes at room temperature. Primary antibodies were diluted in PBSTB and incubated with the samples overnight at 4 °C. After five washes in PBST, samples were incubated with Alexa Fluor-conjugated secondary antibodies (Invitrogen) diluted in PBSTB for 1.5–2.5 hours at room temperature, followed by five additional PBST washes. Samples were mounted using VECTASHIELD mounting medium with or without DAPI (Vector Laboratories; H-1200 or H-1000, respectively). Experimental and control samples were processed on the same slide to allow for direct comparison. The following primary antibodies were used: mouse anti-Armadillo (1:50; DSHB, N27A1) and mouse anti-Prospero (1:20; DSHB, MR1A).

### Image acquisition and processing

Confocal fluorescence images were acquired using a CREST V3 spinning disk confocal system mounted on a Nikon Ti2 inverted microscope, equipped with a 60× Plan Apo 1.4 NA oil immersion objective. Image acquisition was controlled using NIS-Elements software (version 5.3). Shown images represent maximum intensity projections of z-stacks encompassing the first epithelial layer. Identical acquisition settings, including laser power, gain, and camera parameters, were used across all experimental and control samples to ensure comparability. Statistical analyses were performed on raw 16-bit images using Fiji (version 2.0). For presentation purposes, contrast and brightness adjustments were applied uniformly across all images from the same experiment.

Quantification of GFP-positive cells was performed using a custom Fiji macro developed by Dr. Damir Krunic (DKFZ Imaging Facility). This macro was applied to maximum intensity z-projections of either the apical epithelial layer or the full epithelial depth of the midgut. For each fluorescence channel, background subtraction was performed followed by Gaussian blur filtering to enhance feature detection. Signal thresholds were defined for each channel, and the “Find Maxima” function was used to detect cell centers. GFP signal was considered positive if it overlapped with a DAPI-defined nuclear center. The percentage of GFP-positive cells was calculated relative to the total number of DAPI-stained nuclei.

### Single cell RNA-sequencing and high-throughput sequencing

For scRNA-seq, adult flies were starved for 3 hours and subsequently transferred to vials containing filter paper soaked with 5% sucrose for 16 hours. Midguts (n = 20 per genotype) were dissected in ice-cold PBS, ensuring careful removal of the hindgut, crop, Malpighian tubules, and proventriculus. Tissues were digested in 1 mg/mL Elastase (Sigma, #E0258) at 25 °C for 45 minutes with shaking at 1000 RPM, and briefly vortexed every 15 minutes to facilitate dissociation. Following digestion, samples were pelleted, resuspended in PBS, and passed sequentially through 40 μm and 20 μm cell strainers. Cell suspensions were then counted, and approximately 20,000 viable cells per sample were used for single-cell RNA sequencing. Library preparation was performed using the 10x Genomics Chromium Single Cell 5’ V2 kit, following the manufacturer’s protocol. Library quality was assessed using an Agilent Bioanalyzer High Sensitivity DNA chip, and concentrations were determined using a Qubit fluorometer. Libraries were multiplexed and sequenced on an Illumina NextSeq 550 platform at the Deep Sequencing Facility, BioQuant, Heidelberg University.

### scPT-seq primer design

Primers for nested PCR were designed such that both primers flank the gRNA target sites, allowing amplicons to contain the cell barcode, UMI and both gRNA target sites. Specifically, the outer primer was biotinylated on the 5’ end and was separated by a linker, while the nested inner primer was designed to be approximately 50-100bp away from sgRNA2 target site, see Supplemental Table 2 for primer sequences.

### Targeted amplification of CRISPR mutations

To amplify the perturbed gene of interest, a nested PCR strategy was employed. In the first round (PCR1), 20 ng of purified cDNA derived from 10x GEM-RT reactions was used as input. Amplification was carried out using 10 µM partial Read1 forward primer, 10 µM biotin-labeled outer gene-specific reverse primer, and 2X KAPA HiFi HotStart ReadyMix. Thermocycling conditions were as follows: 95 °C for 3 min; 10 cycles of 98 °C for 20 s, 67 °C for 30 s, and 72 °C for 60 s; followed by a final extension at 72 °C for 5 min. PCR1 products were purified using 1.0x volume of AMPure XP beads (Beckman Coulter) according to the manufacturer’s protocol. Biotinylated amplicons were captured using Dynabeads M-270 Streptavidin (Thermo Fisher Scientific), and eluted in 10 mM EDTA (pH 8.2) containing 95% formamide, as per the manufacturer’s instructions, and purified using 1.8x volume of AMPure XP beads. The eluted product was used as input for the second round of PCR (PCR2), performed with 10 µM Partial Read1 forward primer, 10 µM inner gene-specific reverse primer, and 2X KAPA HiFi HotStart ReadyMix. Cycling conditions were identical to PCR, except 15 cycles was performed at 98 °C. PCR2 products were subjected to a second round of Dynabeads M-270 Streptavidin capture and subsequently purified using a 1.8x volume of AMPure XP beads. PCR reactions were done in duplicates and pooled to ensure enough material was available for sequencing.

### Long-read PacBio sequencing

Amplicon libraries were prepared using the PacBio SMRTbell prep kit 3.0 with the following modifications. The initial cleanup step was omitted. During adaptor ligation and cleanup, a 1.8x bead-to-sample ratio was used instead of 1.3x. Nuclease treatment was performed for 7 min (rather than 15 min), followed by cleanup with a 1.8x ratio. The protocol was followed only through the nuclease treatment step. Annealing, binding, and cleanup (ABC) reactions were performed using the SMRT Link–generated ABC protocol, with reagent volumes automatically adjusted to each sample’s molarity. Sequencing was carried out with the Sequel IIe system using the 3.1 binding kit, loading between 130-150 pM where possible (otherwise all material was loaded), and a 30 h movie was acquired.

### Processing of Illumina Reads

Illumina sequencing of the standard 10X reads from the control, *LamC*, and *Prosα3* samples recovered 97M, 85M, and 104M reads respectively for the first replicate and 94M, 96M, and 79M for the second. These were preprocessed into filtered count matrices using CellRanger 7.0.0^38^. Further preprocessing was performed using scanpy 1.10.4^39^. Cells with fewer than 200 genes or with mitochondrial reads above a sample-specific threshold between 9 and 15% were removed. Genes present in fewer than 3 cells were removed. For clustering and annotation, read counts were 1e4 normalized, log1p transformed, scaled to unit variance, and batch corrected using highly variable genes and the mnnpy package^40^. The neighborhood graph was produced with 10 neighbors and 40 PCA components, and leiden clustering performed at a superresolution setting of 2. Cluster cell types were then annotated using marker genes given in Supplemental Table 1.

### Preprocessing of PacBio Reads

PacBio HiFi reads were constructed via Circular Consensus Sequencing (CCS) using the PacBio ccs package 6.3.0, extracted with extracthifi 1.0, and converted to fastq files with samtools 1.21^41–43^. Cell barcodes were identified using the freediv10Xcellbcs package with max error 1 and reject delta 1^44,45^. Reads were mapped to the dm6 BDGP6.28 genome using minimap2 2.22-r1101^46^.

### Background splicing analysis

To assess which effects were due specifically to CRISPR perturbation, we analyzed the natural missplicing rate in the control samples. We conservatively accepted as splice junctions any deletions observed in at least 2 UMIs in the same single cell in the control sample. We identified 14 and 29 natural missplicing junctions in *LamC* and *Prosα3* respectively. See Supp. Fig. 1.3.

### Full Haplotyping

For full haplotyping, we first identified high confidence, high frequency segregating sites in each sample. These were identified directly in the data as belonging to >10% of reads and the major allele being in fewer than 80% of reads covering that site. In both *LamC* and *Prosα3*, and in both the perturbed and control samples, we identified at least one segregating site present in >95% of reads, enabling robust haplotyping of the data. Though clarifying which haplotype is maternal and paternal is not necessary to the downstream analysis, it was identifiable here and included for completeness. The maternal fly line of perturbed and control samples was shared, while the paternal lines were different to provide distinct gRNAs or lack thereof to each sample. Consistent base pairs at each segregating site across control and perturbed samples were thus used to identify the maternal, and therefore also the paternal, haplotype in each sample.

### Mutation calling

The mutation calling pipeline is divided into three parts, corresponding to the level of resolution: Bulk, individual read, and cell-haplotype, that is, all reads from a given chromosome.

Bulk level: Splicing sites and segregating sites are identified as described above.

Read level: The barcode is given from the barcode identification process above. The haplotype is identified from the segregating site in the read closest to target 1. All differences between the reference sequence and the aligned read close to a target site are identified as putative mutations, including all sequencing or other errors but excluding splicing junctions identified from the control sample. An error is considered close to a target site if the closest part of the error is within 10 bp of the cutsite.

Cell-haplotype (chromosome) level: We gather all reads with the same cell barcode and the same haplotype and look at each putative mutation in any read in the set. For each present mutation, we identify the coverage of reads covering that mutation and the number of reads containing that mutation. The quotient gives the mutation read fraction. The mutation with the maximum read fraction is labelled the defining mutation, and the mutation read fraction of the defining mutation is labelled the max frac. The chromosome is then labeled as WT, mutated, or unclassified. Chromosomes with ≥ 10 reads and ≤ 30% max frac are labelled WT. Chromosomes with ≥ 30% max frac representing at least 3 reads are identified as mutated. Chromosomes with insufficient reads to meet these thresholds are unclassified. In mutated chromosomes, reads not containing the defining mutation are removed and all mutations present in > 50% of remaining reads are called, removing the possibility of inconsistent mutation calls on the same chromosome.

Splice junction mutations are identified in three steps. First, they appear as large deletions. Second, splice site modifications are identified as deletions where the left end is a standard left end of a splice junction or the right end is a standard right end of a splice junction, and the other end is more than half the distance to the other standard splice end. Third, the second step is repeated using newly identified splice junction ends as known splice junction ends to allow for mutations to both ends. The third step is repeated until no new splicing junction ends are found. For each canonical splice junction in each chromosome, if a mutation of that splice junction is not found and that splice junction has coverage of at least 3 and is present in fewer than 5% of covering reads, it is labelled as intron retention.

### Mutation abbreviations

Mutations for substitutions are written with the standard [reference bp][position][observed bp] notation, e.g. A12345T. Deletions are abbreviated as D[start position]-[end position], e.g. D12345-12367. Insertions are abbreviated I[insertion position][inserted bases], e.g. I12345ACAATG. Inserted bases are inserted immediately before the insertion position. Splice junction modifications are abbreviated J[standard start position]_[standard end position]>[start position delta]_[end position delta], e.g. J12345_12367>-18_+0 indicates an observed splice junction from 12327 to 12367. Zeros are written with plus signs for clarity. For intron retention, the right half is simply “>-”, e.g. J12345-12367>-. Intron retention is in general a rate rather than a binary quantity, and we are able to quantify this rate. For these two samples, however, we are able to simplify the presentation to intron retention mutation because all splice junctions are present in the overwhelming majority of reads in the control samples, as determined in the background splicing analysis.

### Modeling clone uniqueness

To model clone uniqueness, we first built a background model of the probability of observing a given combination of edits or non-edits at each of the four target sites (2 chromosomes, 2 per chromosome) from the frequency of edits at each target site independently. A combination of mutations at a single target site on a single chromosome is considered one edit here. Fully wt cells were omitted, as we wished to calculate the probability of an observed combination of edits given a cell was edited. As we wished to model the underlying frequency of edits before clonal expansion, we deduplicated cells with the same edits at all four target sites. We furthermore added Laplace smoothing: all single substitutions, insertions up to 3 bp, and deletions. Substitution and insertion smoothing was limited to the range of positions of observed substitutions and insertions, and deletion smoothing to being within 5 bp of observed deletions. One special case had to be modeled separately as it broke the assumption of target site independence: large deletions from one target site to the other. The probability of deletion between the two target sites was thus modeled first, and then the probability of the end positions and surrounding mutations at either end modelled as independent. This background model gives the probability of a clone 𝑐 arising from an independent editing event, 𝑝*_c_*. Let 𝑛 be the number of clones in the sample, and let the number of independent editing events causing clone 𝑐 be given by 𝐸*_c_*. Then assuming the number of clones is large,

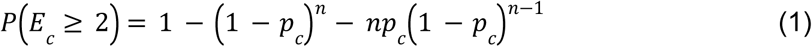

We calculated this for each clone in our data, and removed clones with probability of appearing from more than a single editing event of >0.5%.

### Differential expression analysis

Differential expression analysis was performed using pyDESeq2 0.5.1^47,48^ for each comparison shown in Fig. 3a for each cell type. For each subset of cells for each comparison, we required a minimum number of 10 cells. We further required for each gene that at least one subset of cells for comparison had 10 or more UMIs. For the global Pearson correlation plot, log fold changes were also calculated using pyDESeq2, with the thresholds for cell count and UMIs both reduced to 3.

### Exomic SNP rate analysis

SNPs were identified in a control scRNA-seq sample in regions with 10x or greater coverage using GATK 4.0.9.0, bedtools 2.29.2, and bcftools 1.10.2^49–52^. The number of identified SNPs and the size of the regions with at least 10x coverage were then used to estimate a SNP rate throughout the exome.

### GO term enrichment analysis

GO terms for each gene were downloaded from FlyMine v53^53^. Enrichment p-values were calculated by Fisher exact test and adjusted using Benjamini-Hochberg FDR correction.

### Structural modelling

Structural predictions were generated using AlphaFold2 Multimer via the Cosmic² web service. Protein sequences were obtained from FlyBase (release FB2025_03), and all α and β subunits of the *Drosophila* proteasome were included in the prediction. Among the predicted models, the structure with the highest confidence score was selected for analysis and presentation. The predicted aligned error (PAE) values for the helix of interest (Prosα3, amino acids 80–97) were consistently above 89.4 for the wild-type complex and above 87.4 across the corresponding residues of the mutant complex, indicating a high degree of reliability in the predicted secondary structures of both models. Structural visualization and analysis were performed using UCSF ChimeraX version 1.9 (release date: 2024-12-11). Inter-residue distances were measured using the integrated Distance function within ChimeraX.

## Data availability

Data for this project is available at the Sequence Read Archive under BioProject ID PRJNA1392698 (https://www.ncbi.nlm.nih.gov/sra/PRJNA1392698).

## Code availability

The scptseq software package and notebook tutorials can be found at https://github.com/hawkjo/scptseq.

## Acknowledgements

We would like to thank F. Port, V. Benes, F. Walter, A. Horlova, J. Gleixner, D. Ibberson, D. Grimm, J. Lohmann and W. Huber and for helpful discussions. We thank D. Petrov for help with Pacbio sequencing and discussion. We thank L. Villacorta for help with Oxford Nanopore sequencing. We would like to thank the Nikon imaging facility at Heidelberg University and the EMBL GeneCore sequencing facility for support. This work was supported by the European Research Council (Synergy Project DECODE under grant agreement no. 810296) (M.B and O.S), state funds approved by the State Parliament of Baden-Württemberg for the Innovation Campus Health + Life Science Alliance Heidelberg Mannheim (J.H.), and a Marie Skłodowska-Curie Individual Fellowship (S.R: 894568).

## Author information

### Contributions

Concept of the method: J.H., S.R. Conceptualization and implementation of wet lab experiments: S.R. Conceptualization and implementation of computational pipeline: J.H. Single-cell RNA-sequencing: S.R., S.L., T.W. Pacbio library preparation and sequencing: M.O., H.O., S.R. Structural modelling: M.H., J.H., S.R. Investigation: J.H., S.R., S.L., T.W., M.B., O.S. Data analysis: J.H., S.R. Supervision: O.S., M.B., S.R. Figure preparation: J.H., S.R., M.H. Writing original draft: S.R, J.H. Writing, reviewing and editing: J.H., S.R., O.S, M.B, with input from all authors.

