## Supplemental Materials for "Direct detection of CRISPR mutations and transcriptional responses at single cell resolution *in vivo*"

*Contributed equally, #Corresponding author

| Method | gDNA based | RNA based | Same cells | High throughput | Whole transcriptome | Substitutions | Insertions | Deletions | Detection of proteins | Splicing change detection | Variant caller haplotyping | Full haplotyping | Missing Chromosomes | Lineage tracing | *In vivo* |
|---|---|---|---|---|---|---|---|---|---|---|---|---|---|---|---|
| scPT-seq | ✗ | ✓ | ✓ | ✓ | ✓ | ✓ | ✓ | ✓ | ✗ | ✓ | ✓ | ✓ | ✓ | ✓ | ✓ |
| TISCC-seq | ✗ | ✓ | ✓ | ✓ | ✓ | ✓ | ✗ | ✗ | ✗ | ✗ | ✗ | ✗ | ✗ | ✗ | ✗ |
| SDR-seq | ✓ | ✗ | ✓ | ✓ | ✗ | ✓ | ✓ | ✓ | ✗ | ✗ | ✓ | ✗ | ✗ | ✗ | ✗ |
| CRAFTseq | ✓ | ✗ | ✓ | ✗ | ✓ | ✓ | ✓ | ✓ | ✓ | ✗ | ✓ | ✗ | ✗ | ✗ | ✗ |
| scSNV-seq | ✓ | ✗ | ✗ | ✓ | ✓ | ✓ | ✗ | ✗ | ✗ | ✗ | ✓ | ✗ | ✗ | ✗ | ✗ |

**Supplemental Figure 1.1: Feature comparison of recent methods with sequencing of single-cell CRISPR edits.**

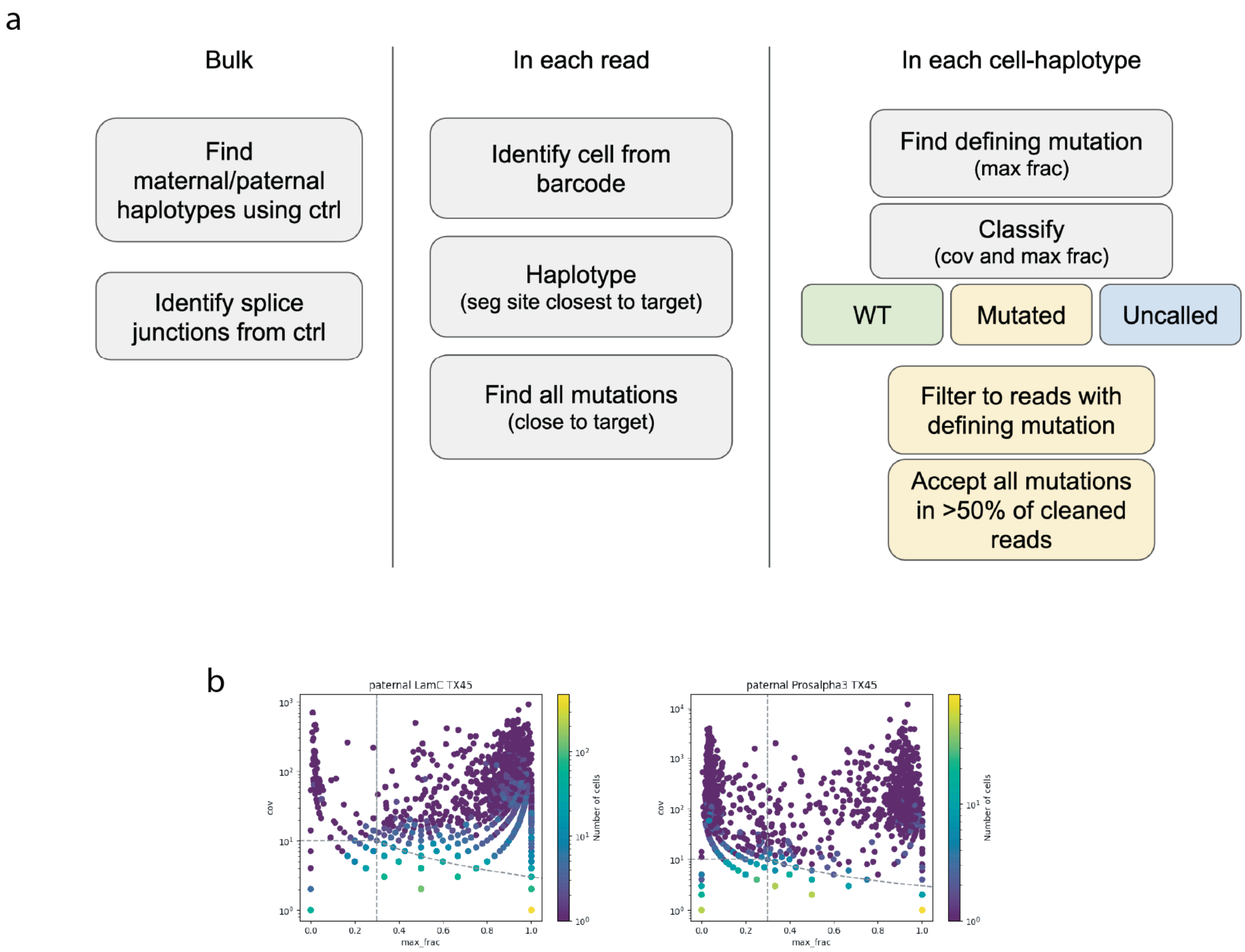


**Supplemental Figure 1.2: Mutation calling algorithm and example data.** a. Overview of algorithm for haplotyping, calling splicing junctions, and identifying mutations. b. Coverage vs. max frac (see Methods) for example data with LamC and Prosalpha3 mutations.

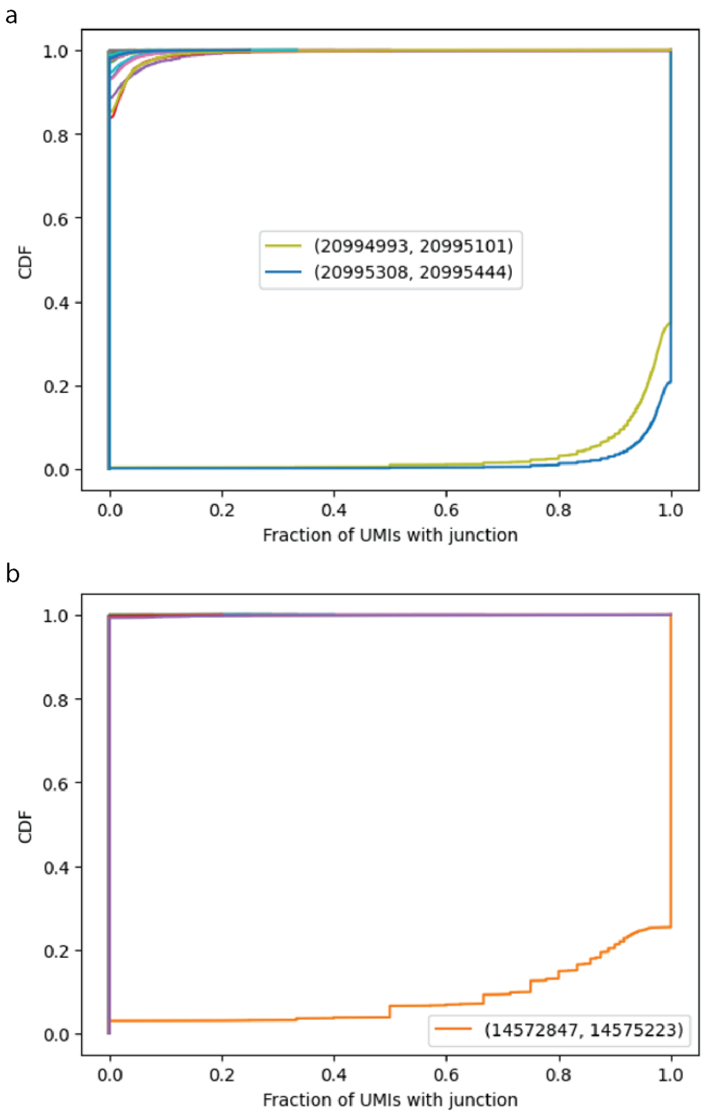


**Supplemental Figure 1.3: Observed splicing junction in control data.** All observed splicing junctions with at least two UMIs in a single cell in a. Prosalpha3, with 29 observed junctions in up to >15% of cells, and b. LamC, with 14 observed junctions in small numbers of cells. CDF over cells. Legends show canonical splicing junctions.

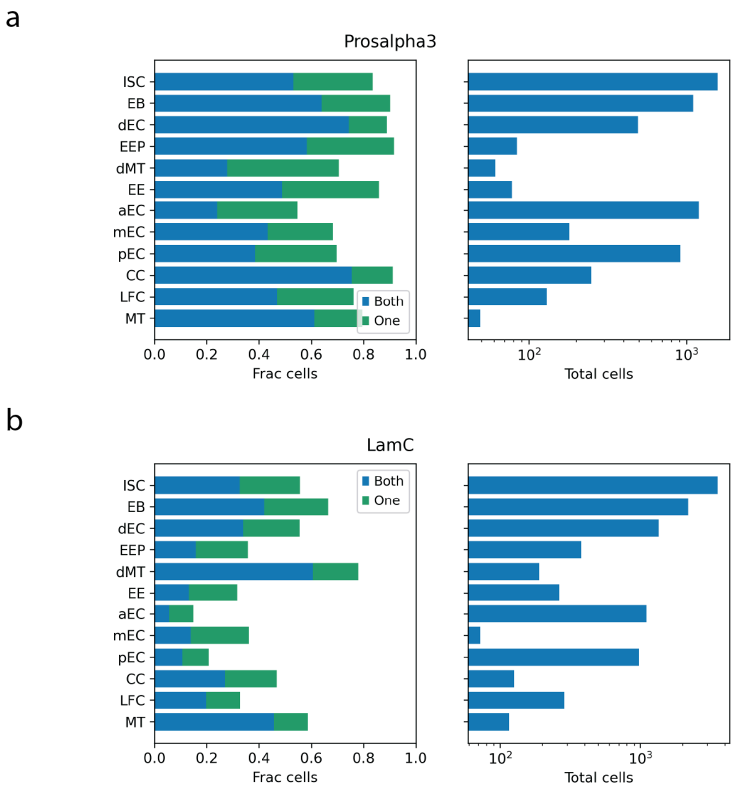


**Supplemental Figure 1.4: Recovery of mutation status across cell types.** For a. Prosalpha3 and b. LamC: Left panel: Fraction of cells in which one or both chromosomes have recovered mutation statuses, stratified by cell type. Right panel: Corresponding number of cells for each cell type.

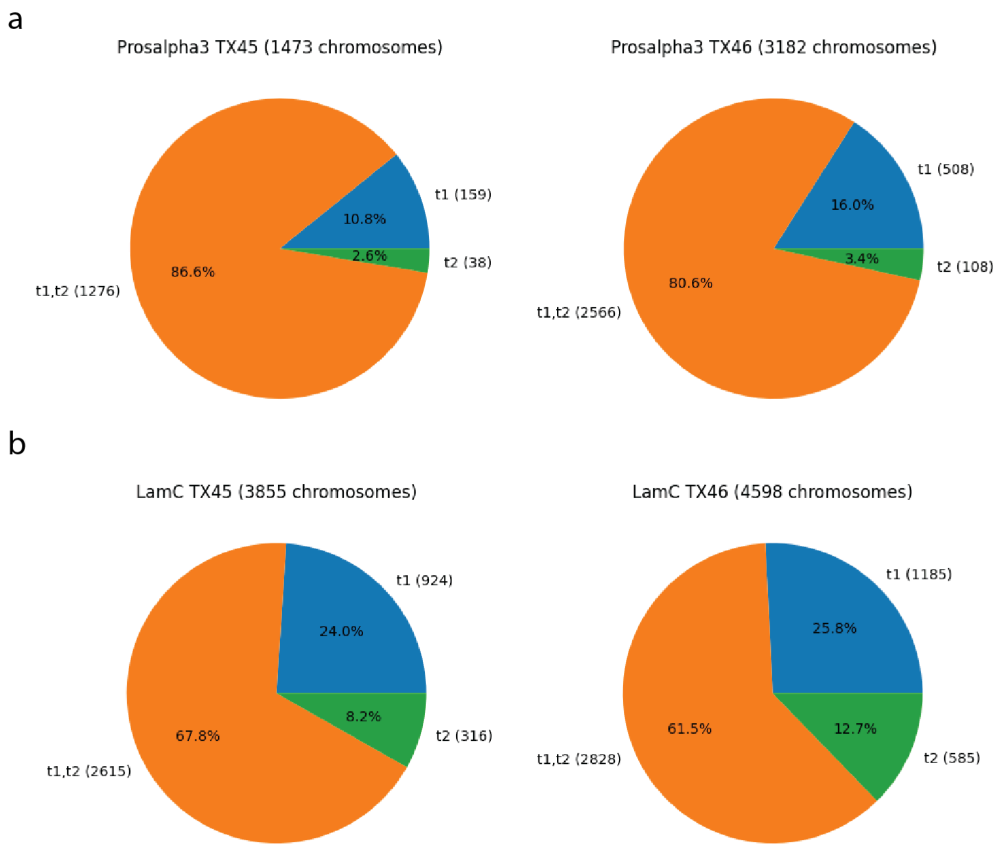


**Supplemental Figure 2.1: Target site specificity.** Target site specificity for Target 1 and Target 2 in a. Prosalpha3 and b. LamC. Between replicates the overall efficiency is slightly different, but the target site preference is highly reproducible.

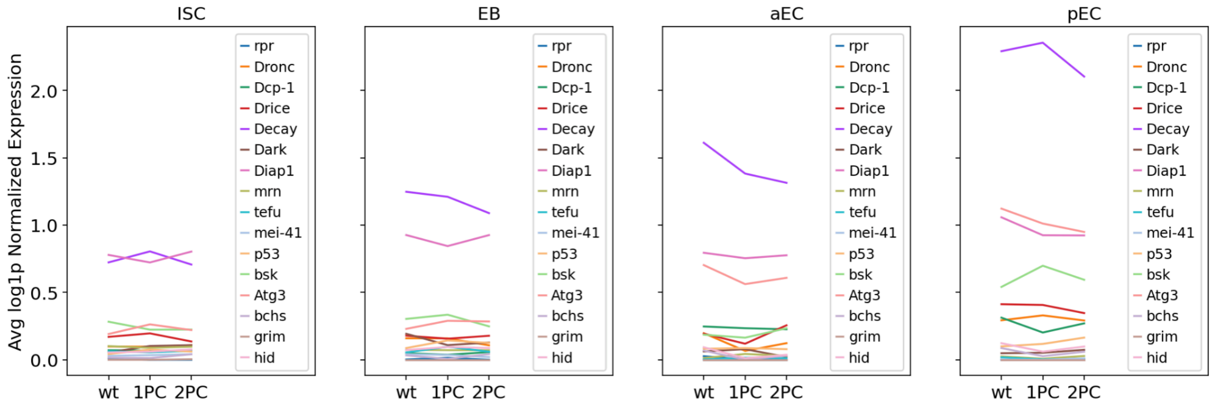


**Supplemental Figure 2.2: Apoptosis marker expression.** Apoptosis marker expression vs. mutation status in *Prosα3* samples in the four most common cell types.

a b

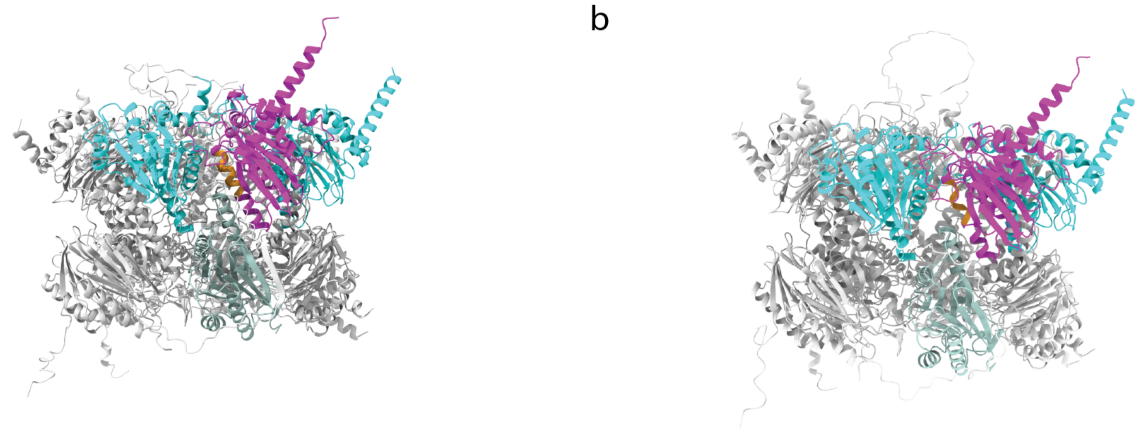

**Supplemental Figure 2.3: AlphaMultimer proteasome core complex structure prediction.** Proteasome core complex structure prediction with a. WT and b. the most common 12 bp deletion in our data.

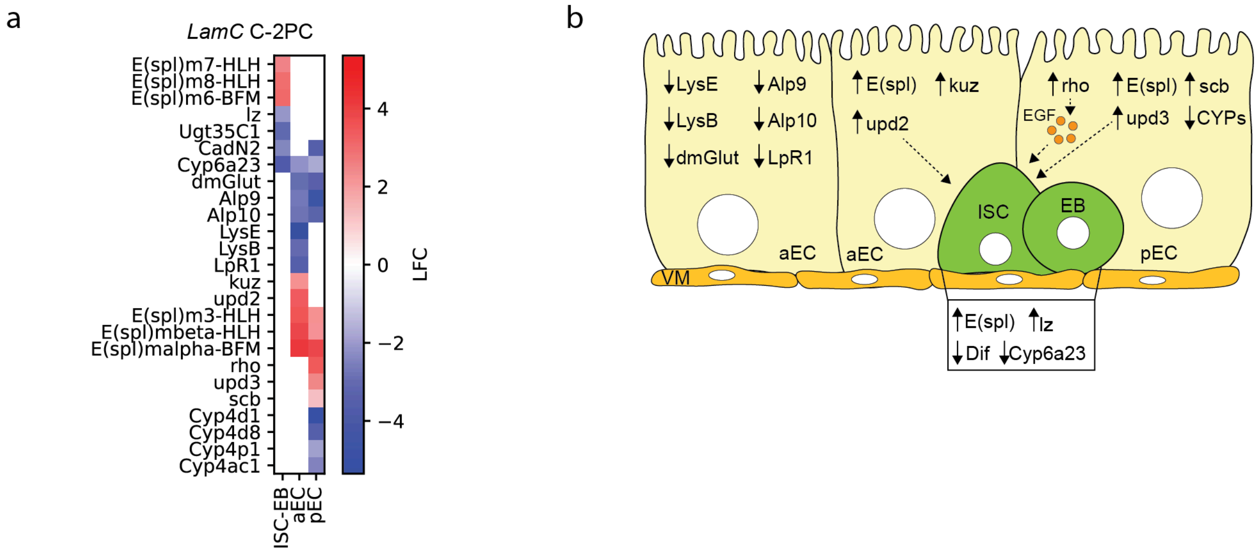


**Supplemental Figure 3.1: LamC DE results. a.** Expression log fold change for genes of notable biological interest in our major cell types. **b.** A model of genes of interest affecting stem cell proliferation.

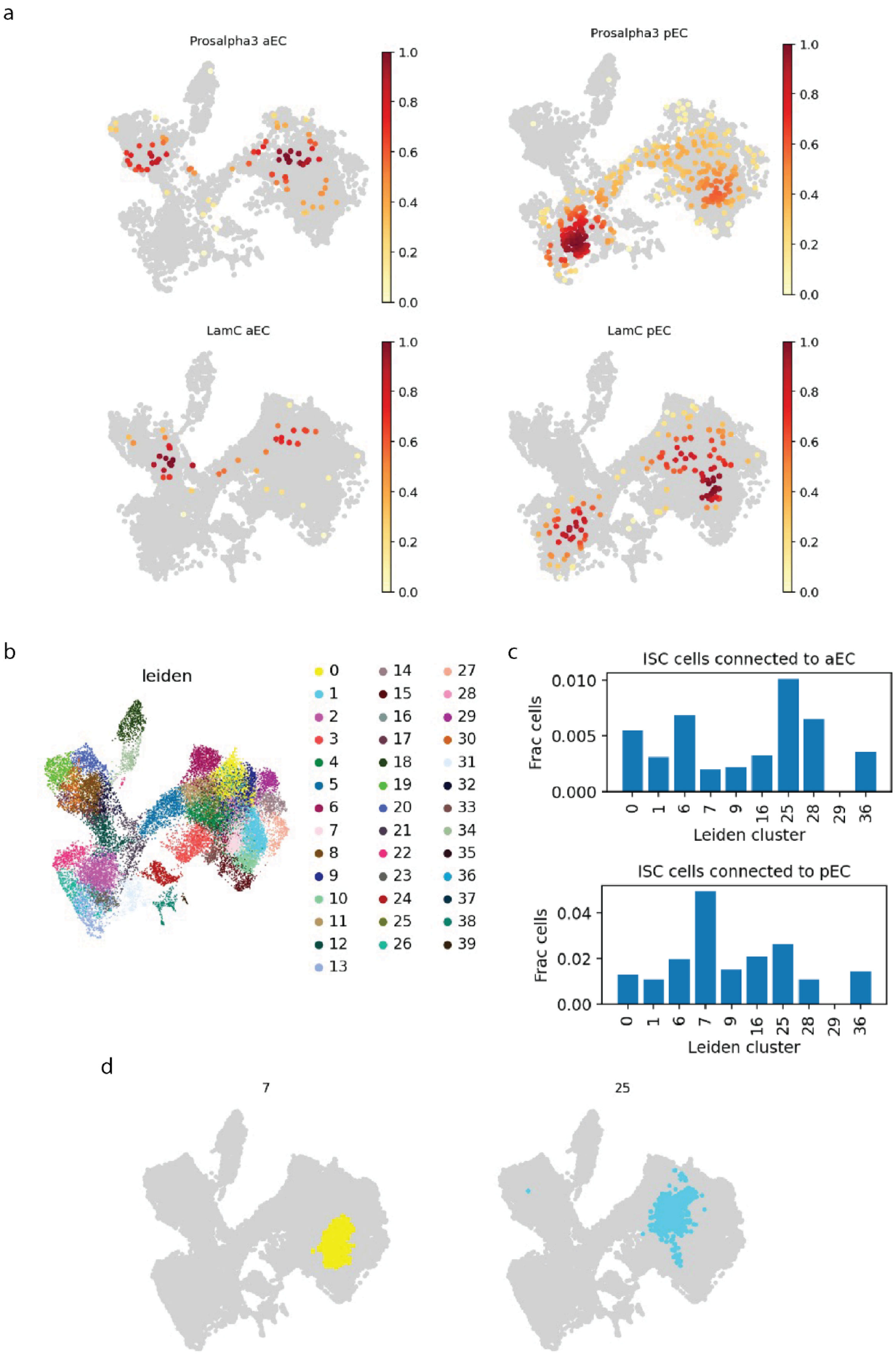
a
Prosalpha3 aEC
Prosalpha3 pEC
LamC aEC
LamC pEC
1.0
0.8
0.6
0.4
0.2
0.0
b
leiden
0 1 2 3 4 5 6 7 8 9 10 11 12 13 14 15 16 17 18 19 20 21 22 23 24 25 26 27 28 29 30 31 32 33 34 35 36 37 38 39
c
ISC cells connected to aEC
Frac cells
0.000
0.005
0.010
0 1 6 7 9 16 25 28 29 36
Leiden cluster
ISC cells connected to pEC
Frac cells
0.00
0.02
0.04
0 1 6 7 9 16 25 28 29 36
Leiden cluster
d
7
25

**Supplemental Figure 4.1: Clonal analysis detail. a.** The data from Figure 4d stratified by perturbation target. **b**. Leiden superclustering of UMAP used to identify subclusters of ISC cells enriched with spatially linked cells. **c.** Enrichment of aEC- and pEC-linked ISCs in each listed cluster. **d.** Leiden clusters 7 and 25, the subclusters most enriched with posterior- and anterior-linked cells respectively.

| Cell type | Marker Genes |
|---|---|
| ISC | Dl, spdo |
| EB | N, Su(H) |
| EB/EEP | esg |
| EEP | hdc |
| EE | pros, 7B2 |
| dEC | nub |
| aEC | alphaTry, betaTry |
| mEC | thetaTry, Npc2f |
| pEC | lambdaTry, LManVI |
| CC | Vha100-4, lab |
| LFC | PGRP-SC1b, Jon65Ai |

**Supplemental Table 1: Marker genes for cell type annotation**

| Primer description | Primer sequence |
|---|---|
| Read 1 | CTACACGACGCTCTTCCGATCT |
| Prosalpha3 nested | TTCCGCCGTACTGAGTGTACGCCTG |
| Prosalpha3 5BiotinTEG | GATCCGACTGGTACAGTTGGTAGCCG |
| LamC nested | GCGAGCTCCTTGGCCTGATCCTC |
| LamC 5BiotinTEG | TGCGAAGATCATCCAGTTGGCGGCG |

**Supplemental Table 2: Primers**
